# Ancient genomes from southern Africa pushes modern human divergence beyond 260,000 years ago

**DOI:** 10.1101/145409

**Authors:** Carina M. Schlebusch, Helena Malmström, Torsten Günther, Per Sjödin, Alexandra Coutinho, Hanna Edlund, Arielle R. Munters, Maryna Steyn, Himla Soodyall, Marlize Lombard, Mattias Jakobsson

## Abstract

Southern Africa is consistently placed as one of the potential regions for the evolution of *Homo sapiens*. To examine the region’s human prehistory prior to the arrival of migrants from East and West Africa or Eurasia in the last 1,700 years, we generated and analyzed genome sequence data from seven ancient individuals from KwaZulu-Natal, South Africa. Three Stone Age hunter-gatherers date to ~2,000 years ago, and we show that they were related to current-day southern San groups such as the Karretjie People. Four Iron Age farmers (300–500 years old) have genetic signatures similar to present day Bantu-speakers. The genome sequence (13x coverage) of a juvenile boy from Ballito Bay, who lived ~2,000 years ago, demonstrates that southern African Stone Age hunter-gatherers were not impacted by recent admixture; however, we estimate that all modern-day Khoekhoe and San groups have been influenced by 9–22% genetic admixture from East African/Eurasian pastoralist groups arriving >1,000 years ago, including the Ju|‘hoansi San, previously thought to have very low levels of admixture. Using traditional and new approaches, we estimate the population divergence time between the Ballito Bay boy and other groups to beyond 260,000 years ago. These estimates dramatically increases the deepest divergence amongst modern humans, coincide with the onset of the Middle Stone Age in sub-Saharan Africa, and coincide with anatomical developments of archaic humans into modern humans as represented in the local fossil record. Cumulatively, cross-disciplinary records increasingly point to southern Africa as a potential (not necessarily exclusive) ‘hot spot’ for the evolution of our species.

## Main text

Archaeological, fossil and genetic data consistently place the earliest traces of anatomically modern humans in sub-Saharan Africa^1–6^. East Africa often features in human origins studies, because the earliest modern human remains, dating to ~190 kya (kya = thousand years old/ago), originates from Ethiopia^2,3^. In southern Africa, cross-disciplinary data however converge, indicating that understanding the population histories of the region can contribute to understanding the origins of our species^4,7–10^.

Recent syntheses indicate the occupation of the southern African landscape by the genus *Homo* from about 2 Mya^11^, with a major transitional phase between 600 kya and 200 kya (from the Earlier Stone Age into the Middle Stone Age)^12^. Current interpretation of the fossil record indicates the presence of archaic *H. sapiens* at >200 kya, during the earlier phases of the Middle Stone Age, and anatomically modern humans from ~120 kya^11^. After ~120 kya we also see some of the earliest archaeological evidence for modern human behavior and thinking in sub-Saharan Africa^13^. Genetic studies (mitochondria, Y-chromosome, and autosomes) constantly identify southern African Khoe-San populations as carrying more unique variants and more divergent lineages than any other living groups^1,8–10,14–17^. The deepest population split among modern humans – between Khoe-San and other groups – is estimated to ~160–100 kya, based on short sequence fragments^15,16^, and genome-wide SNP data^8^. Some of these patterns were used to argue for a southern African origin of humans^10^, although others suggested several regions^8,18^.

Middle Stone Age sites in KwaZulu-Natal, South Africa, e.g., Sibudu Cave^13^, Umhlatuzana Rock Shelter^19^ and Border Cave^20^, demonstrate human occupation since >100 kya. Here we report the sequencing and analyses of the genomes of seven ancient individuals from KwaZulu-Natal (Table 1, Figure 1), directly dated to the last 2 kya. The genomes of individuals from Ballito Bay and Doonside represent the first genomic data for Stone Age hunter-gathers prior to the arrival of migrants from within and outside of Africa <2 kya. These data reveal previously unknown admixture patterns for southern African indigenous groups, and push the emergence of modern humans back to >260 kya.

**Table 1.**
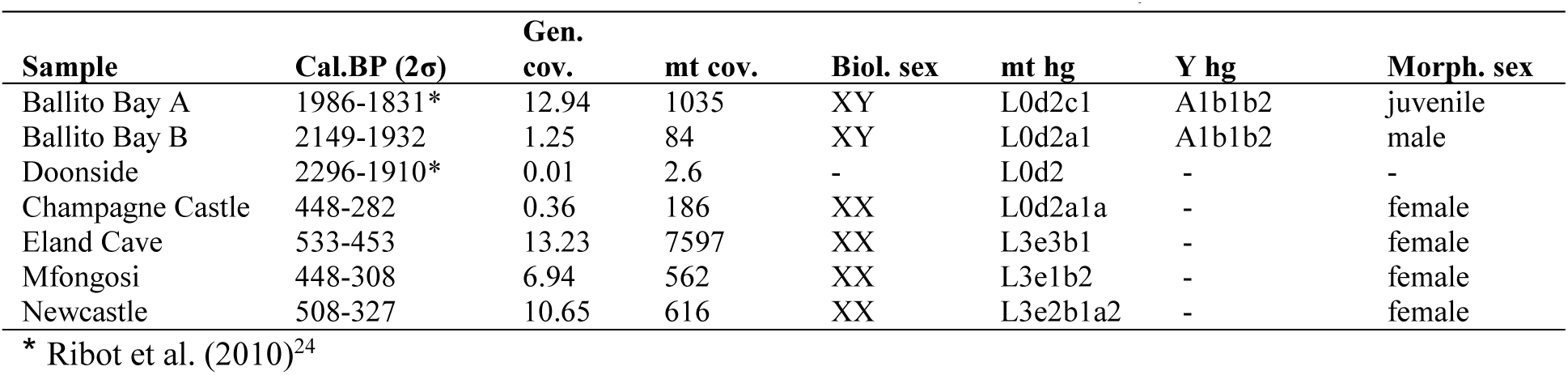
Sample information. Summary table of calibrated dates, genomic and mitochondrial DNA coverage, sex estimations and mitochondrial and Y-chromosomal haplogroup assignments (see SI section 1–5 for further information). All new radiocarbon dates are reported in SI section 1.

**Figure 1.**
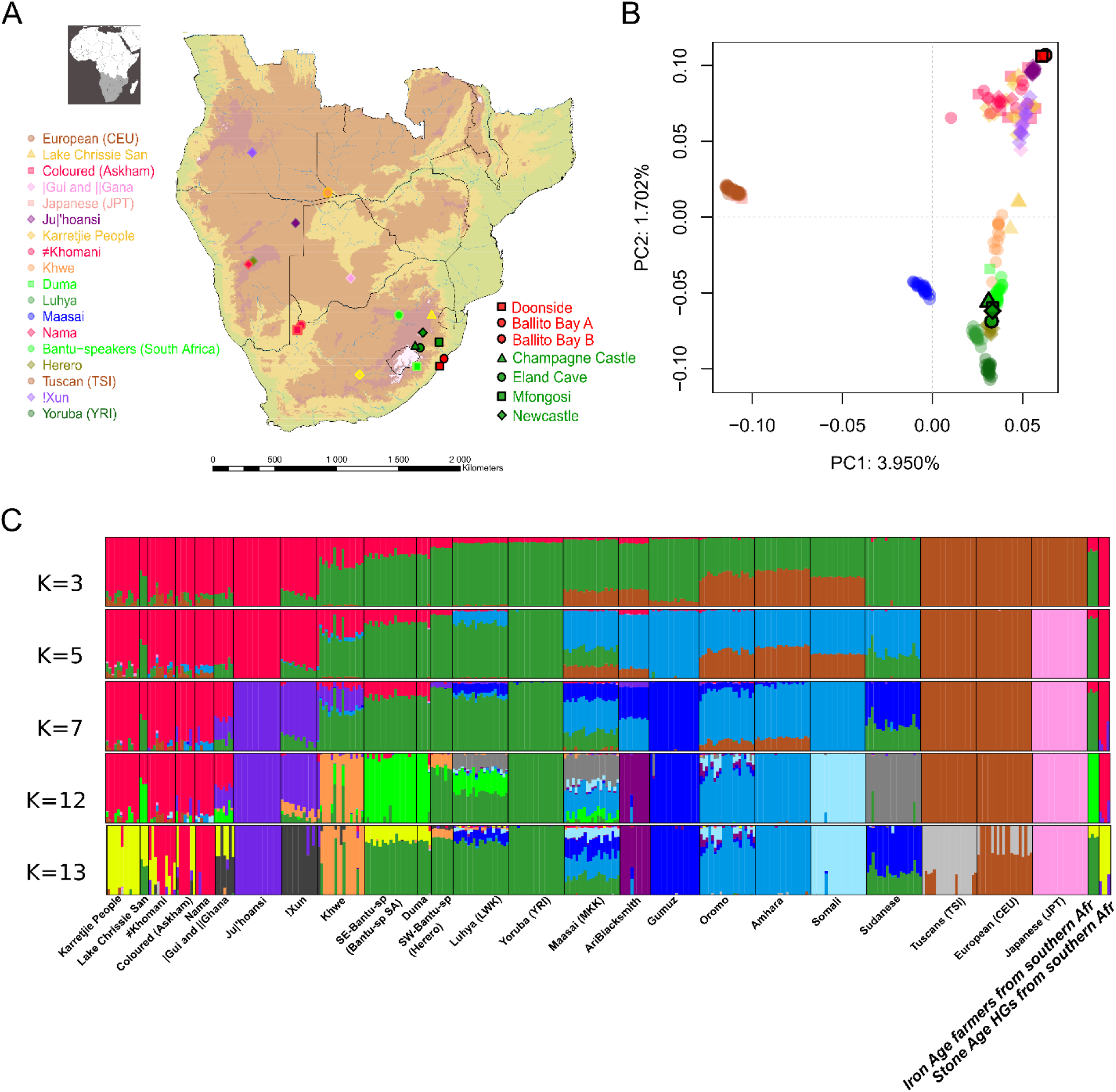
(A) Locations of the archaeological sites and the geographic centers of comparative populations from Schlebusch et al^8^. (B) PCA with southern African, African and global comparative data. (C) Admixture analysis, displayed for selected number of clusters (K), see figure S6.5 for all K values.

We sequenced the genomes of three Stone Age hunter-gatherers and four Iron Age farmers, directly radiocarbon dated to ~2 kya and 0.5–0.3 kya respectively, to between 0.01x and 13.2x genome coverage (Figure 1, Table 1, SI 1 for archaeological contexts, and SI 2–3 for sampling and laboratory procedures). The DNA-sequence data display all features characteristic of ancient DNA (e.g.^21^, short DNA fragments and consistently higher cytosine deamination at fragment ends, SI 4). Five individuals were morphologically sex-assigned (SI 1), and have now been genetically confirmed (SI 5). The previously unknown sex of the Ballito Bay A juvenile has been genetically determined as male (Table 1, SI 1, 5).

The three Stone Age individuals, Ballito Bay A Ballito Bay B, and Doonside, and an Iron Age individual from Champagne Castle, carry mitochondrial sub-haplogroups belonging to haplogroup L0d (SI 5.2), common in current-day Khoe-San populations^14^. The remaining three Iron Age individuals, from Newcastle, Eland Cave, and Mfongosi, have mtDNA haplogroups that fall within L3e, common to current-day Bantu-speaking groups^14^. Both males from Ballito Bay carry the Y chromosome A1b1b2 haplotype (SI Section 5.1), common among modern-day Khoe-San^17^.

To assess population affinities among the ancient individuals and their relations to modern-day groups, we merged the ancient southern African genome data with published genotype datasets from southern Africa^8,9^, Africa as a whole, and from across the globe (SI Section 6, Table S6.1). We further merged and investigated the ancient genome data with a set of complete genomes of 11 individuals from across the world^22^, including individuals from southern, eastern and western Africa (SI Section 6). Principal Component Analysis (PCA) and admixture analyses show that the three Stone Age individuals are related to present-day Khoe-San groups, specifically to southern Khoe-San, such as the Karretjie People^8^ and the Lake Chrissie San^23^ (Figs. 1B-C, Extended Data Figs. 1, 7, 8, Figs. S6.1-S6.5, S9.6-S9.9). The four Iron Age individuals all group with populations of West African origin/descent, specifically with southeast Bantu speakers from South Africa (Figs. 1B-C, S6.1-S6.6). This observation is consistent with archaeological evidence for the arrival of migrant Iron Age farmers of West African descent to the eastern parts of southern Africa at ~1.7 kya^24^.

The Stone Age individuals form one extreme in the PCA (that separates Khoe-San individuals from all other Africans and non-Africans, Fig. 1B and S6.2). All modern-day Khoe-San are drawn towards other Africans and non-Africans compared to the ancient individuals from Ballito Bay, including Ju|’hoansi San, thus far thought to be the least affected by recent admixture^8,9^. A more detailed PCA points towards Eurasian and/or east African admixture into all modern-day Khoe-San (Fig. 1B, S6.3, Extended Data Fig. 1). We tested various admixture-scenarios^25^ into Khoe-San groups using the good-coverage, high quality, genome of Ballito Bay A (SI 6.4–6.5) and conclude that the admixture source was an (already admixed) Eurasian/East African group (Extended Data Fig. 2, Extended Data Table 1 and 2), comparable to the Amhara, today living in Ethiopia (SI Section 6.4–6.5). We estimate that Ju|’hoansi individuals received 9–14% (Extended Data Fig. 2, Extended Data Table 2, SI 6.4–6.5) admixture from this mixed East African/Eurasian group (69%/31%, Extended Data Fig. 2, and SI Section 6.5), and all Khoe-San groups show 9–22% of this admixture (Fig. 2, Extended Data Table 2, SI 6.4–6.5).

**Figure 2.**
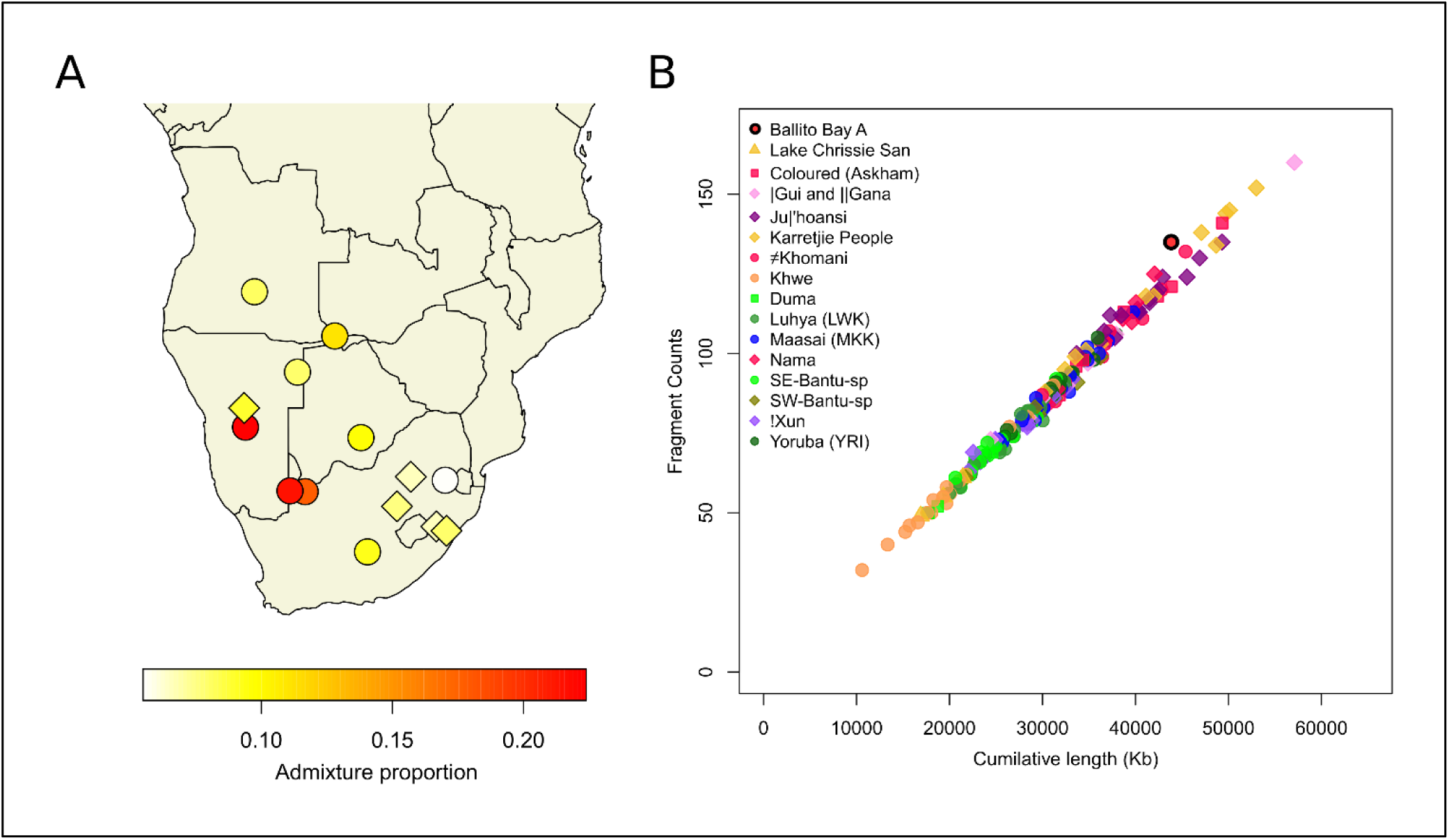
(A) East African/Eurasian admixture proportions (inferred with f_4_ ratio test using Amhara) in a southern African comparative dataset^8^. Circles are San and Khoekhoe populations and diamonds are Bantu-speakers. (B) Runs of Homozygosity of the 200–500 Kb bin in Africans (see fig S7.2 for inclusion of non-Africans).

We dated this admixture event into the Khoe-San using admixture LD decay patterns^25^ to between 50 (±3) generations ago for the Ju|’hoansi (San) and 44 (±4) generations for the Nama (Khoekhoe) corresponding to 1.5–1.3 kya (assuming 30 years/generation) (SI 6.6). The East African/Eurasian source of the admixture is particularly pronounced in herding Khoe groups such as the Nama (Extended Data Table 2, SI 6.4–6.5). Based on these results, we suggest a migration from East Africa into southern Africa, resulting in admixture with local hunter-gatherers ≥1.5 kya. This scenario is consistent with a model of herding practices being introduced from the northeast by migrating pastoralists^8,26,27^. The migration had a pronounced impact on all current Khoe-San groups, not only on the descendants of Stone Age herders, such as the Khoekhoe (Fig. 2B, Extended Data Table 2, SI 6.4–6.5). This has been an elusive result since all modern-day Khoe-San individuals display ≥9% recent admixture. The East African/Eurasian admixture into San and Khoekhoe groups resulted in elevated diversity in present-day Khoe-San groups (Fig 3, Extended Data Fig. 3, SI Section S7.1-S7.2). The admixture also inflates inference of past effective population sizes^28,29^, where modern-day San individuals display elevated effective population sizes compared to other African groups (including Ballito Bay A, Extended Data Fig. 4, SI 7.3).

**Figure 3.**
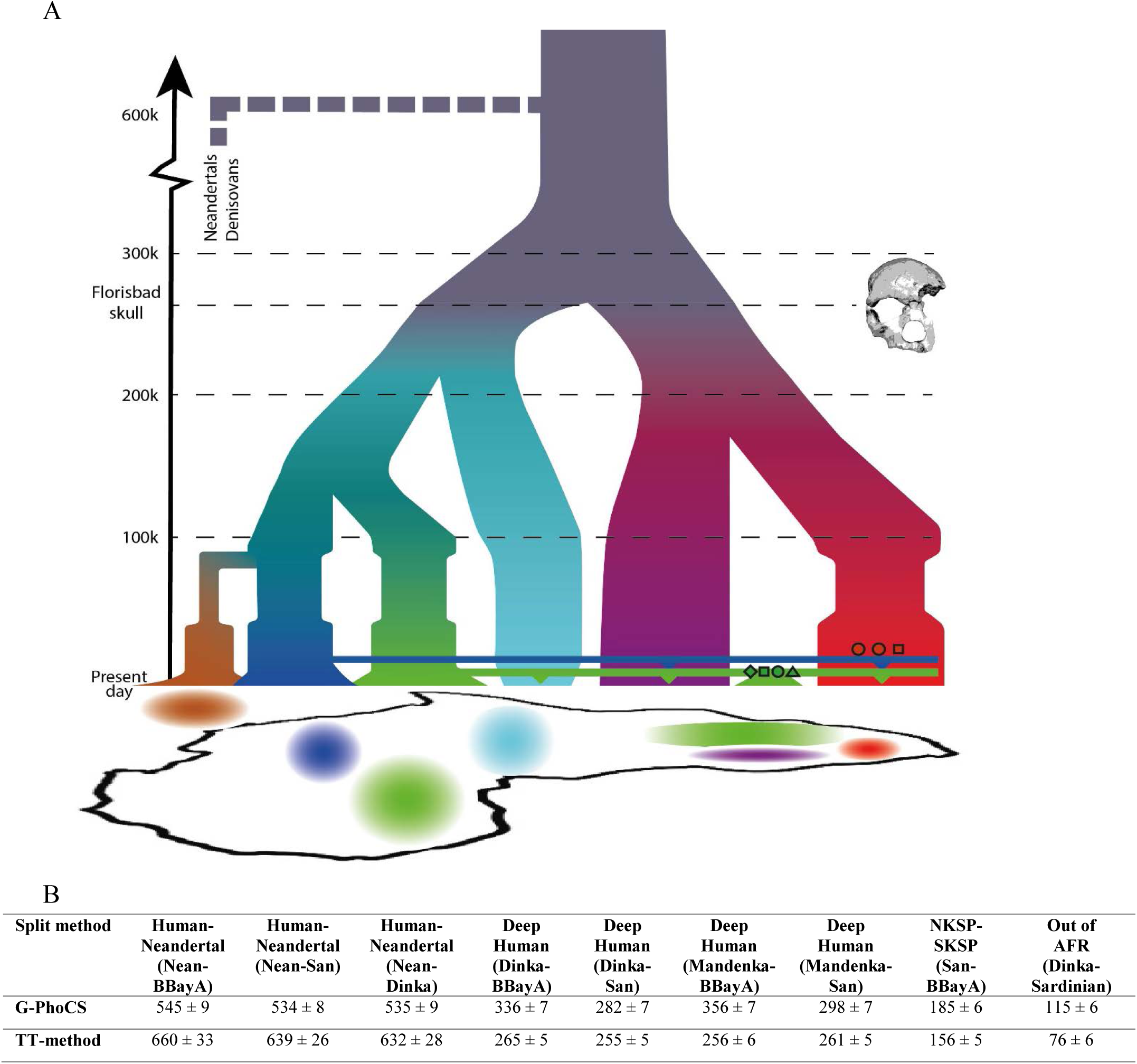
Demographic model of African history and estimated divergences. (A) The estimated age of the Florisbad skull from southern Africa is indicated by a dashed line. The ancient southern African individuals in this study are denoted using the labels in Fig 1A, where the Stone Age hunter-gatherers are shown by red symbols, and Iron Age farmers of West African origin as green symbols. The depiction of population split-times, hierarchy and population sizes (width along a horizontal axis for populations) are a summary of the results presented in Extended Data Figures 1–2, 4–6. (B) Estimated population split times using the Gronau et al. approach (G-PhoCS)^15^ and the novel Two-by-Two (TT) method described in SI 9. The deepest divergence among humans is estimated to 260–360 kya. Divergence time estimates for human vs Neandertal, and non-Africans vs Africans (indicating the out of Africa event) are given for reference, and these estimates are consistent with recent estimates in the literature^4^. NKSP-SKSP shows the estimated split time between Ballito Bay A and Ju|’hoansi.

To decipher early human history, we used several complementary approaches^15,25,28^, and developed a novel coalescent-based approach (the various demographic events discussed are depicted in Figure 3). We focus on the good coverage, high-quality ancient genomes, in particular the Stone Age hunter-gatherer boy, Ballito Bay A. In contrast with modern-day southern African individuals, he was unaffected by admixture with herders from East Africa^8,26,27^, Bantu-speaking farmers from West Africa, or Eurasian immigrants. Specifically, we estimated divergences between various sets of individuals based on diploid called sites of Ballito Bay A (SI Section 8) and 12 previously published high coverage modern and archaic human genomes^22,30^ using a coalescent-based approach (G-PhoCS,^15^, SI 8), assuming 1.5×10^−8^ mutations/generation^31^ and 30 years/generation. We estimate the split times between Ballito Bay A and modern-day individuals (excluding Ju|’hoansi) to 285–356 kya (Fig 3, Extended Data Fig. 5, SI 8), and the deepest split time to 356±7 kya ago for the comparison with the Mandenka of West Africa (Extended Data Fig. 5). Population split times using the admixed Ju|’hoansi instead of Ballito Bay A are on average 55,000 years younger (average: 268 kya based on Ju|’hoansi *vs* average: 323 kya based on Ballito Bay A, Extended Data Fig. 5, Table S8.1). This difference is likely caused by Ju|’hoansi carrying 9–14% admixture from East Africans/Eurasians, and may also be impacted by intrinsic properties of ancient DNA^21^(SI section 4).

We developed a novel method (SI section 9) based on the ‘concordance’ approach^8,32^ that alleviates assumptions about past population sizes, and is robust to low levels of admixture. Briefly, assuming a general split model without migration, and picking 2 chromosomes (from two different individuals or from one individual with diploid data) from each of two subpopulations, it is possible to analytically derive the parameters of the model based on the frequencies of the 8 possible polymorphic sample configurations (assuming an infinite sites model and a known ancestral variant state). The method estimates the population split-time separately for each branch in a two-population model, resulting in two estimates of the same split. It provides the possibility to estimate, independently, the split between the Stone Age Ballito Bay A boy and other groups, using genetic data from modern-day individuals avoiding bias caused by properties of ancient DNA^21^, and the need for phased data. We evaluate this approach, demonstrating that split-time estimates are accurate, little affected by low levels of admixture/migration (SI section 9.1), and improve with genome coverage (SI section 9.1). For the Ballito Bay A *vs* Dinka split, the two branches are estimated to 301±5 kya (Ballito Bay A) and 265±5 kya (Dinka, Fig 3b, Extended Data Fig. 6, SI 9.1), demonstrating that the deepest split among humans is >260 kya; even if we base the analysis on the genetic variation in the Dinka to avoid possible impact of ancient DNA properties (SI section 9). The Ju|’hoansi *vs* Dinka split has similar split times for the two estimated branches (258±5 kya, Ju|’hoansi and 255±5 kya, Dinka, Fig 3b Extended Data Fig. 6 and SI section 9.1), thus some 43–10 kya less than the Ballito Bay A *vs* Dinka split. This difference is likely due to the additional admixture from East Africans/Eurasians into present-day Khoe-San reported above (SI 6).

Cumulatively, our data show that the deepest split among modern humans occurred at >260 kya (Figure 3), pushing the emergence of *H. sapiens* to beyond 260 kya. Potential additional gene-flow between southern African hunter-gatherers and other groups >2 kya would only lead to the estimates provided here being underestimates of the true population split time. This deep divergence at >260 kya is halfway to the human *vs* Neandertal/Denisovan split (Fig 3, SI 8–9, Table S9.1,^30^), and as deep as the split between the Denisovans and Neandertals^4,30^. The deepest split among humans is also 2.5–3.5 times as deep as human migration out of Africa, and predates the next oldest split in human population history by ~100 kya (African rainforest hunter-gatherers *vs* western/eastern Africans). We acknowledge that mutation-rate estimates are debated, varying based on methodology. Recently, a consensus rate of 1.5×10^−8^ per base pair per generation has emerged^31^, but this estimate might also be revised, affecting the chronological dating of events inferred by genomic data. Thus, although our assumptions on mutation rates and generation times influence the exact chronological estimates, our results notably increase the time depth for the deepest split for modern humans on a relative scale.

Several studies point to the possibility of deep population structure among sub-Saharan groups from central and West Africa^4,33–35^, but not for the Khoe-San. It is possible that some fraction of the deep split times between Ballito Bay A (and Ju|’hoansi if restricted to modern-day individuals) and other modern-day sub-Saharan individuals/groups can be explained by low levels of deep structure/admixture. But, unless it is common to all non-San groups, it is unlikely to have a substantial effect on the split time estimates in this study. Interestingly, West African populations partly capture this deep population structure (Extended Data Fig 2), but it is not seen in East African groups (SI 6.5).

The San, often represented by Ju|’hoansi, has consistently been included in influential investigations on human evolutionary history as a ‘non-admixed’ population^4,6,8,10,16,22,30^ to date the deepest splits among modern humans, to infer admixture patterns of archaic humans outside of Africa, and to study the population history of sub-Saharan Africa. Many of these inferences may be biased by the recent admixture into all San groups.

For the reconstruction of robust population histories and origins, evidence from the archaeological, fossil and genetic records should ideally converge ^7^. The successful genetic sequencing of the seven ancient individuals from KwaZulu-Natal contributes to the better understanding of two relatively recent events that are clear in the archaeological record. One is the migration of East African herders introducing pastoralism ≥1.5 kya to southern Africa, and their mixing with local hunter-gatherers. The other is the large-scale migration of Iron Age Bantu-speaking farmers from West Africa introducing agriculture in southern Africa. We have established, for the first time, a new pre-admixture genetic baseline for hunter-gatherers from southern Africa, based on the three oldest individuals. The genetic data for these ~2 kya individuals from KwaZulu-Natal provide a record of the people on the landscape at the time, and inform on the deep human history represented by hunter-gatherers of this region.

The outcome of the deep divergence time for *H. sapiens,* evidenced through the Ballito Bay A boy, requires integration with the local archaeological and fossil records. The deepest split-time calculation of >260 kya is consistent with the archaeological estimate for the onset of the Middle Stone Age across sub-Saharan Africa^12^. Southern African archaeological sites with absolute dates >160 kya are still rare, e.g., Cunene in Namibia, Border Cave, Florisbad, Kathu Pan 1, Pinnacle Point Cave 13B and Wonderwerk Cave in South Africa^13^. Dated Middle Stone Age contexts of ~300–250 kya are thus far limited to three layers at Florisbad^36^, and one at Kathu Pan 1^37^. At the Florisbad site in central South Africa, Middle Stone Age artefacts were found together with human remains dating to 259±35 kya^36^. Amongst the remains is a partial cranium, with a cranial capacity similar to that of modern humans, interpreted as representing a combination of archaic and modern characteristics^36,38^. Human remains from Hoedjiespunt, South Africa of ~300–200 kya were ascribed to *H. heidelbergensis*, because although morphologically modern, they seemed larger than modern-day Africans^39^. These records attest to the presence of humans on the southern African landscape at the time of the earliest modern human divergence, predating 260 kya, and the fossils deserve closer morphological scrutiny. Whether the Florisbad skull represents a modern human ancestor, or an archaic form of human who contributed little or no genetic material to modern humans is an open question, as is our potential relationship with *Homo naledi* who also roamed the South African landscape between about 236 kya and 335 kya^40^. Although we do not rule out that the ancestors of KwaZulu-Natal Stone Age hunter-gatherers might have originated elsewhere in sub-Saharan Africa, or might have mixed with other groups >2 kya, we suggest that archaeological, fossil and genetic records increasingly point towards a modern human development that includes southern Africa.

## Materials and methods

See supplementary information for full description of methods and analyses.

## Acknowledgements

We thank Dr. Carolyn Thorpe, Dr. Gavin Whitelaw and Mudzunga Munzhedzi at the Kwa-Zulu Natal Museum for access to the museum and Luciana Simões and Mário Vicente for technical assistance. This project was supported by grants from Knut and Alice Wallenberg foundation (to MJ), the Swedish Research council (no. 642–2013-8019 MJ, and no. 621–2014-5211 to CS), the Göran Gustafsson foundation (to MJ), a Wenner-Gren foundations postdoctoral fellowship (to TG), and African Origins Platform grant from the South African National Research Foundation (to ML). Sequencing was performed at the National Genomics Infrastructure (NGI), Uppsala, and computations were performed at Uppsala Multidisciplinary Center for Advanced Computational Science (UPPMAX). Data is available for download through the European Nucleotide Archive (EBI-ENA) under project xxx.

## Author contributions

C.M.S., H.M., M.L. and M.J. conceived the study, M.L. coordinated local stakeholders and permitting, M.S. conducted morphological analysis. H.M., A.C. and H.E. sampled the material and performed DNA laboratory work, C.M.S, H.M., T.G., P.S., A.C, A.R.M. and M.J. analyzed the genomic data, M.L. provided fossil and archaeological interpretations, C.M.S., H.S. and M.L. provided historical and ethnographic interpretations, and C.M.S, M.L. and M.J. wrote the paper with input from all authors.

## Extended data

**Extended Data Fig. 1:**
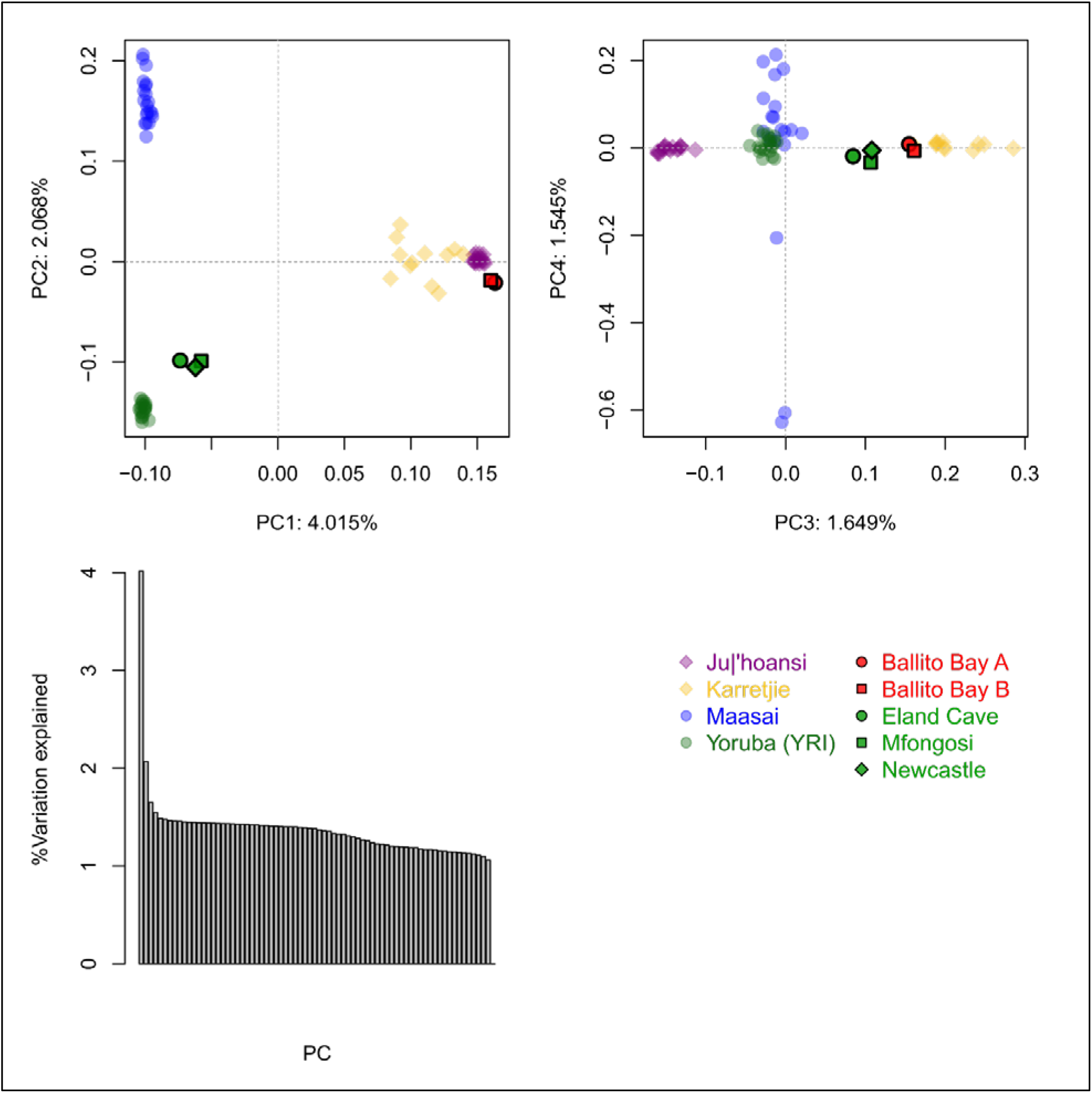
Principal Component analysis with comparative East and West Africans (Maasai and Yoruba) and southern and northern Khoe-San (Karretjie People and Ju|’hoansi), excluding Champagne Castle and Doonside to maximize the number of retained SNPs. (A) Ju|’hoansi is shifted towards Maasai (east Africans) compared to Ballito Bay A for PC 1 and 2, while some of the Karretjie individuals are shifted towards Maasai and some towards Yoruba (PC1 and 2). (B) Ballito Bay A and Ballito Bay B cluster with Karretjie People (southern San) and not with Ju’|hoansi (northern San) for greater PCs. The Khoe-San admixture in the Iron Age farmers also cluster with the Karretjie People (C) Bar plot displaying the amount of genetic variation explain by each PC.

**Extended Data Table 1:**
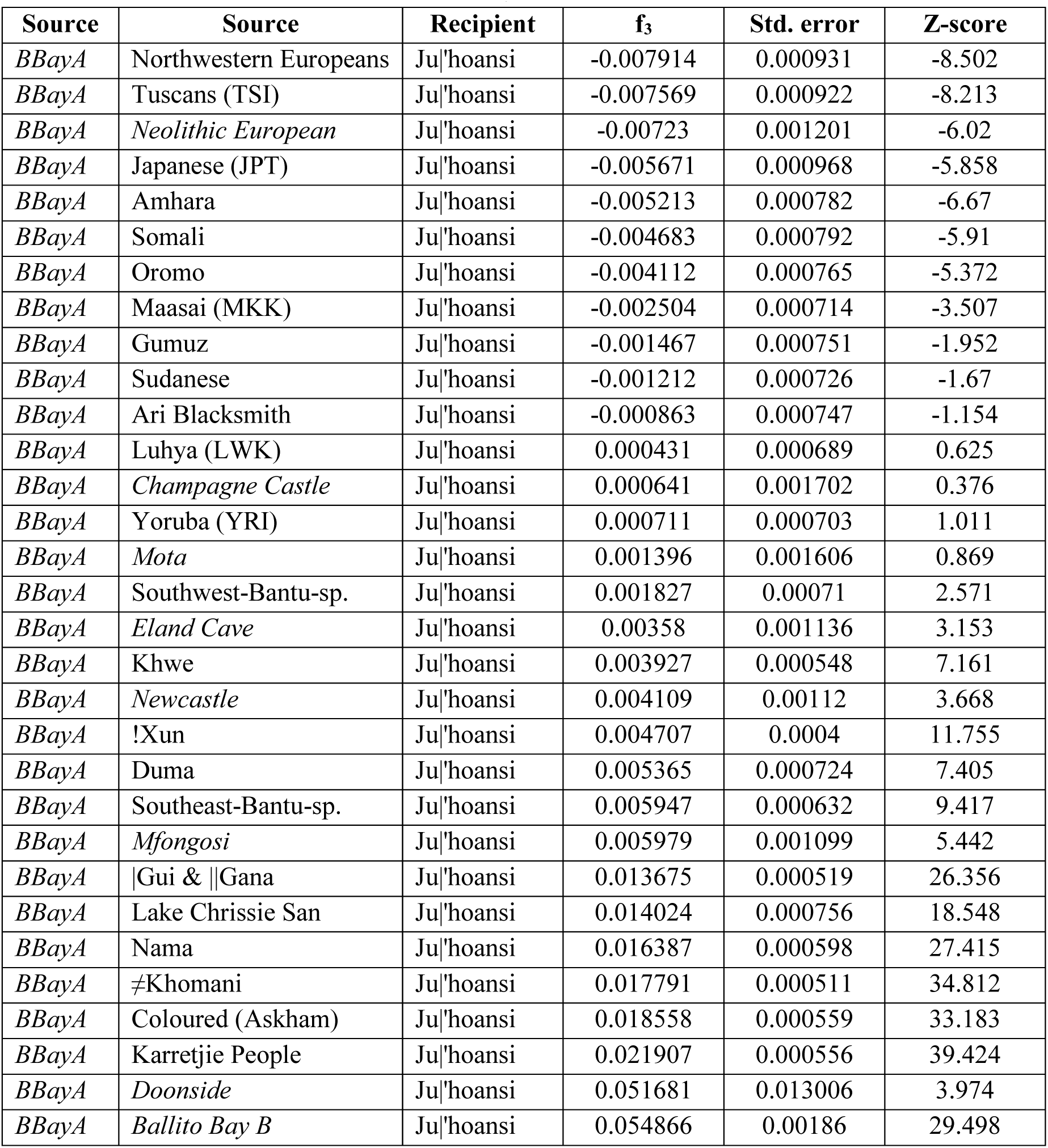
Admixture into Ju|’hoansi inferred using f_3_ statistics. To estimate whether the Ju|’hoansi received admixture from another population, we computed the f_3_ statistic^25^ with Ju|’hoansi as the recipient population, the diploid Ballito Bay A as one source and other populations of the East African extended dataset as the other source population. Negative Z scores was observed for all non-Africans and East Africans (except the prehistoric Mota individual), as a source population in addition to Ballito Bay A. This shows admixture from either East Africans, Eurasians, or a mixture of the two into the Ju|’hoansi. Ancient individuals are marked in italic.

**Extended Data Fig. 2:**
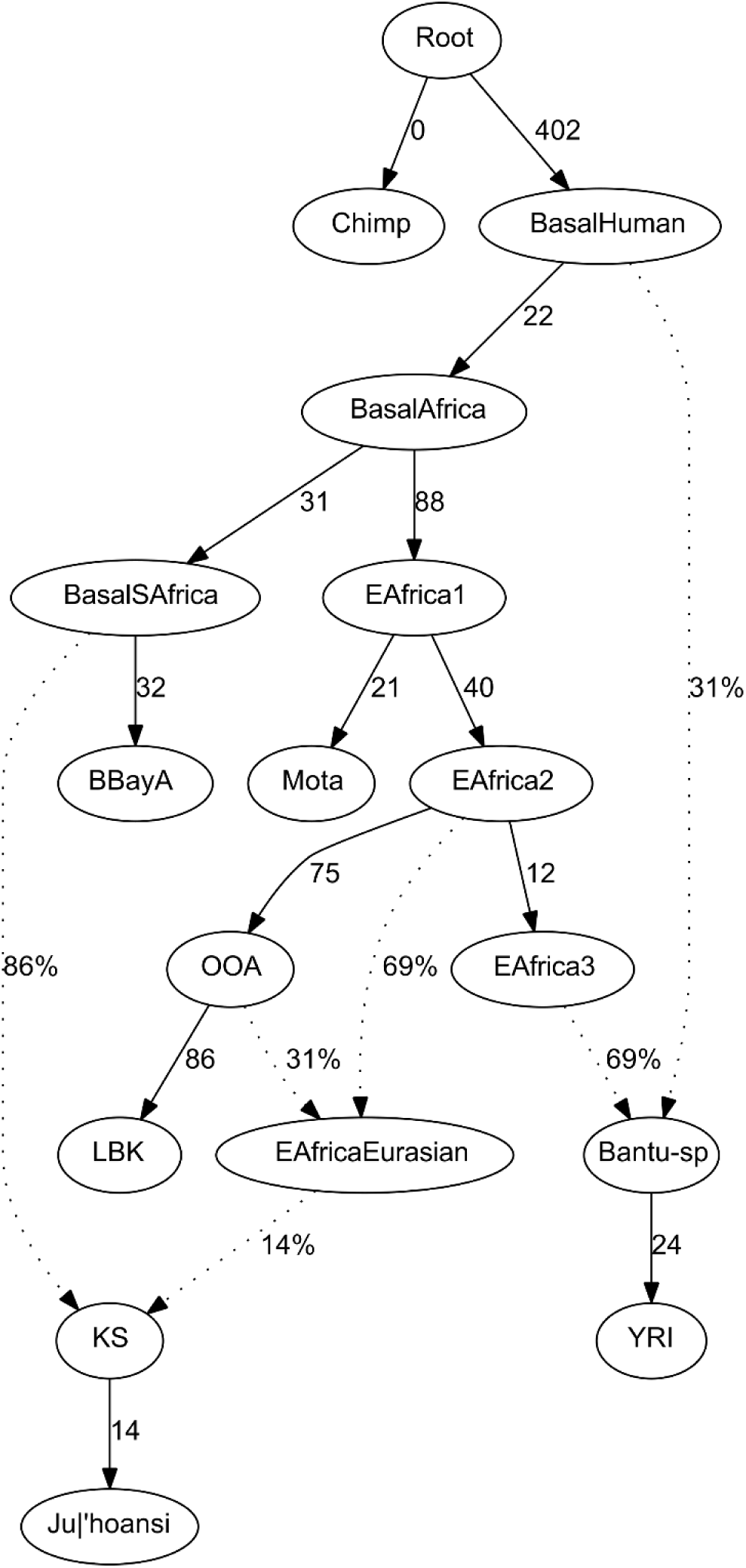
A population and admixture graph model of Ju|’hoansi as an admixed population between southern Africans and an admixed (Eurasian/East African) population is consistent with the data. The numbers next to edges represent the amount of drift between the nodes (multiplied by 1000). The model is including Yoruba as a potential source of Bantu-speaking ancestry. Ju|’hoansi as a Khoe-San population without admixture from Bantu-speakers is also consistent with the data. Alternative tested models constructed in a hierarchical way are discussed in SI section 6.5. The most likely model for the Ju|hoansi is shown here.

**Extended Data Table 2:**
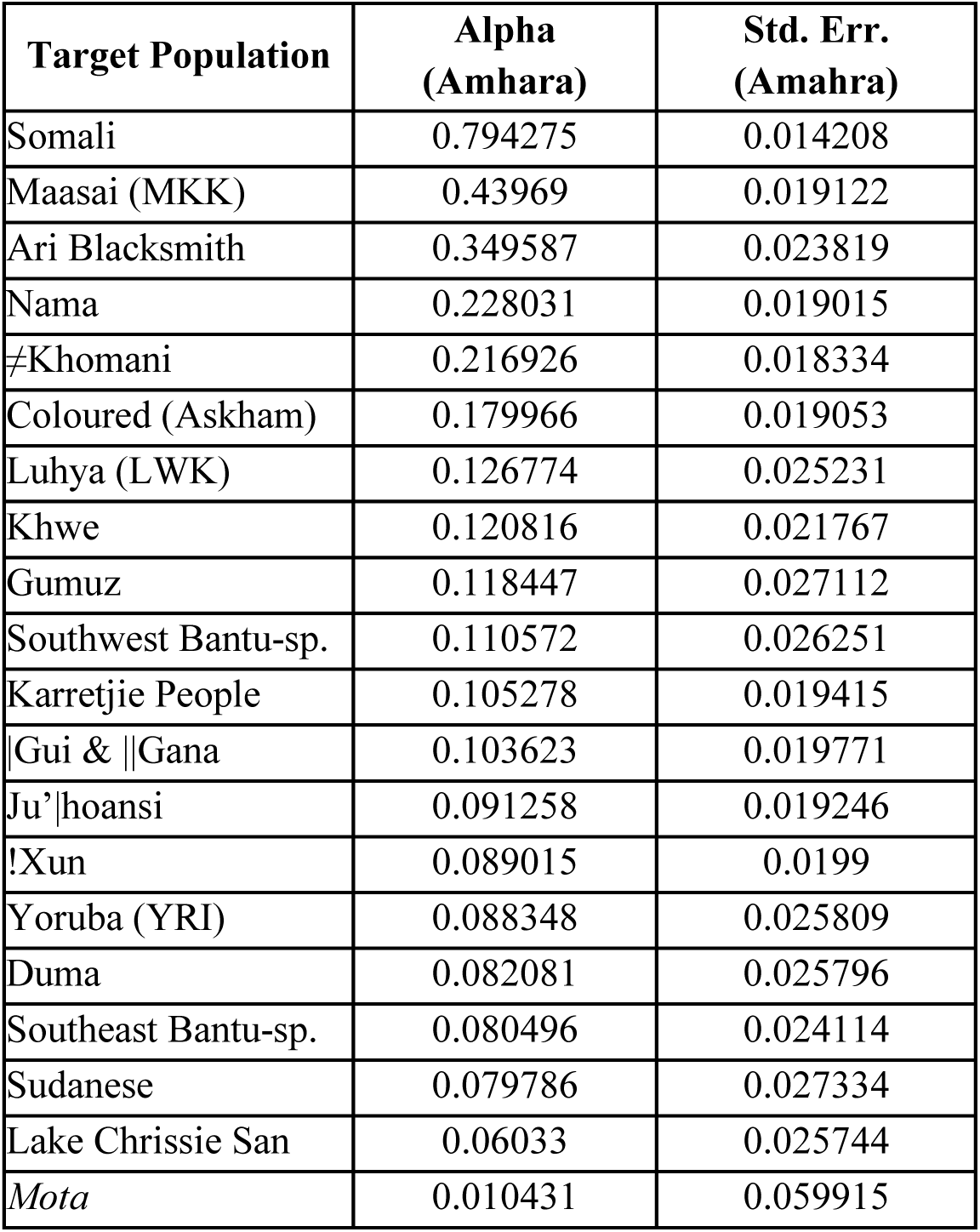
f_4_ ratios estimating East African/Eurasian ancestry proportions in Africans from the East African extended dataset. As southern African populations contain admixture from a source of mixed East African and Eurasian ancestry, we used f_4_ ratios to estimate this ancestry using a modern-day east African group (Amhara) that display Eurasian admixture as source. We use the Amhara as a proxy for admixing source. Other sources from the same and other datasets are shown in SI section 6.4. The ancient individual is marked in italic.

**Extended Data Fig. 3:**
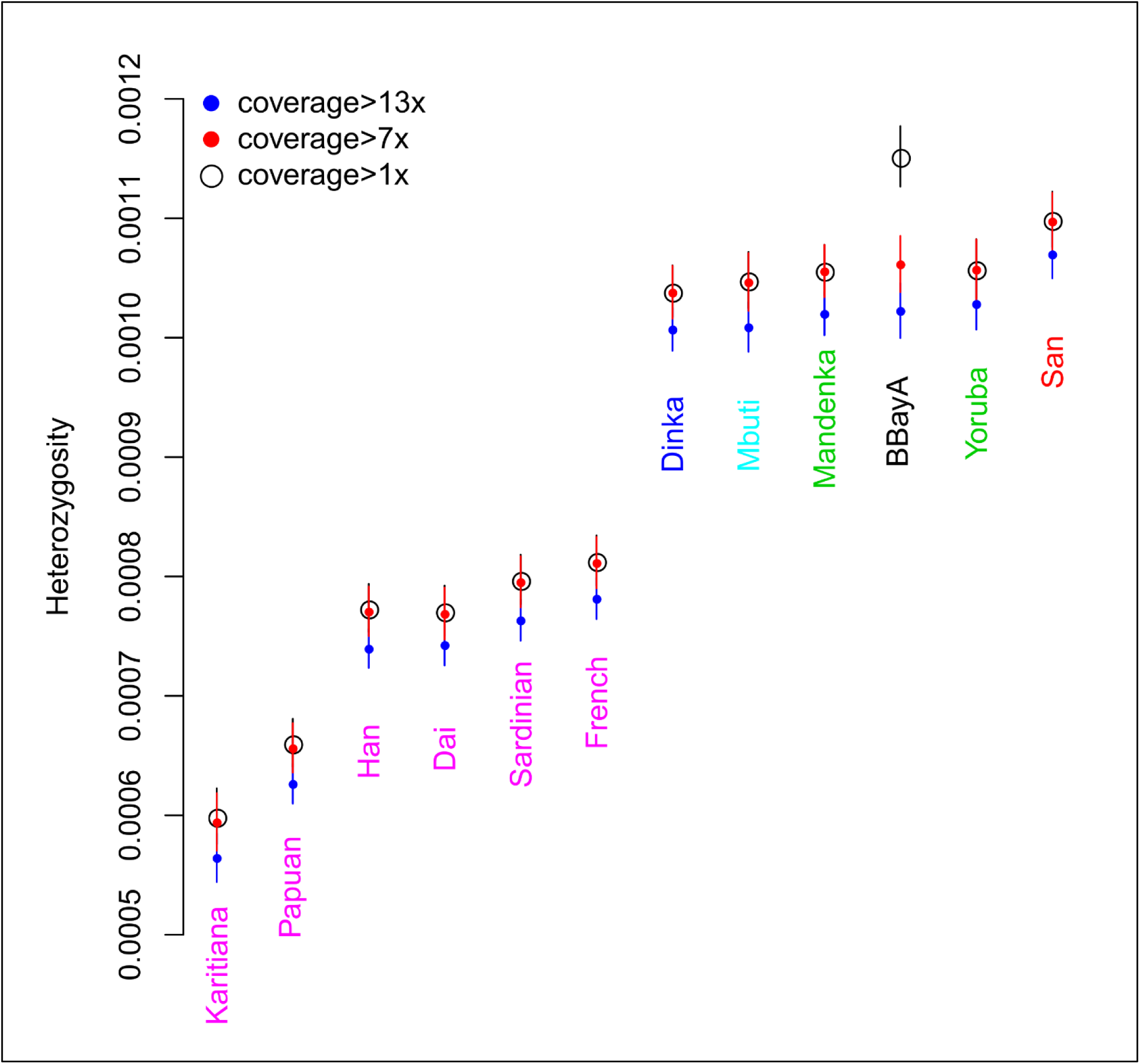
Heterozygosity estimates based on stringently filtered sites. Red points indicates heterozygosity based on all sites with coverage >7x, black circles on sites with coverage >1x and blue points have additional applied quality filters, including >13x (see SI section 7.1). Points are sorted in ascending order based on the more stringently filtered sites (blue points).

**Extended Data Fig. 4:**
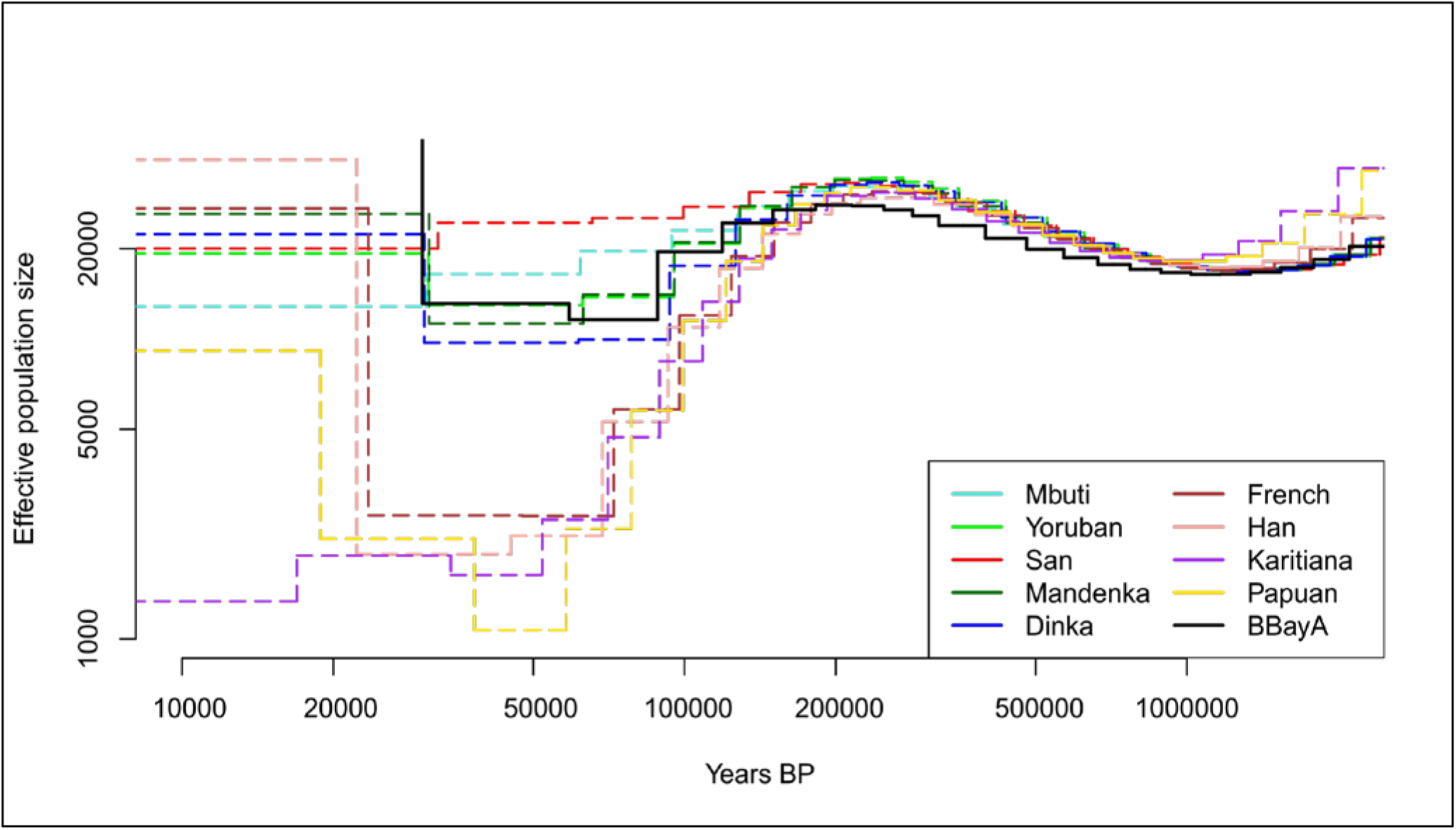
MSMC plot of 11 HGDP genomes together with the diploid full genome of Ballito Bay A (BBayA).

**Extended Data Fig. 5:**
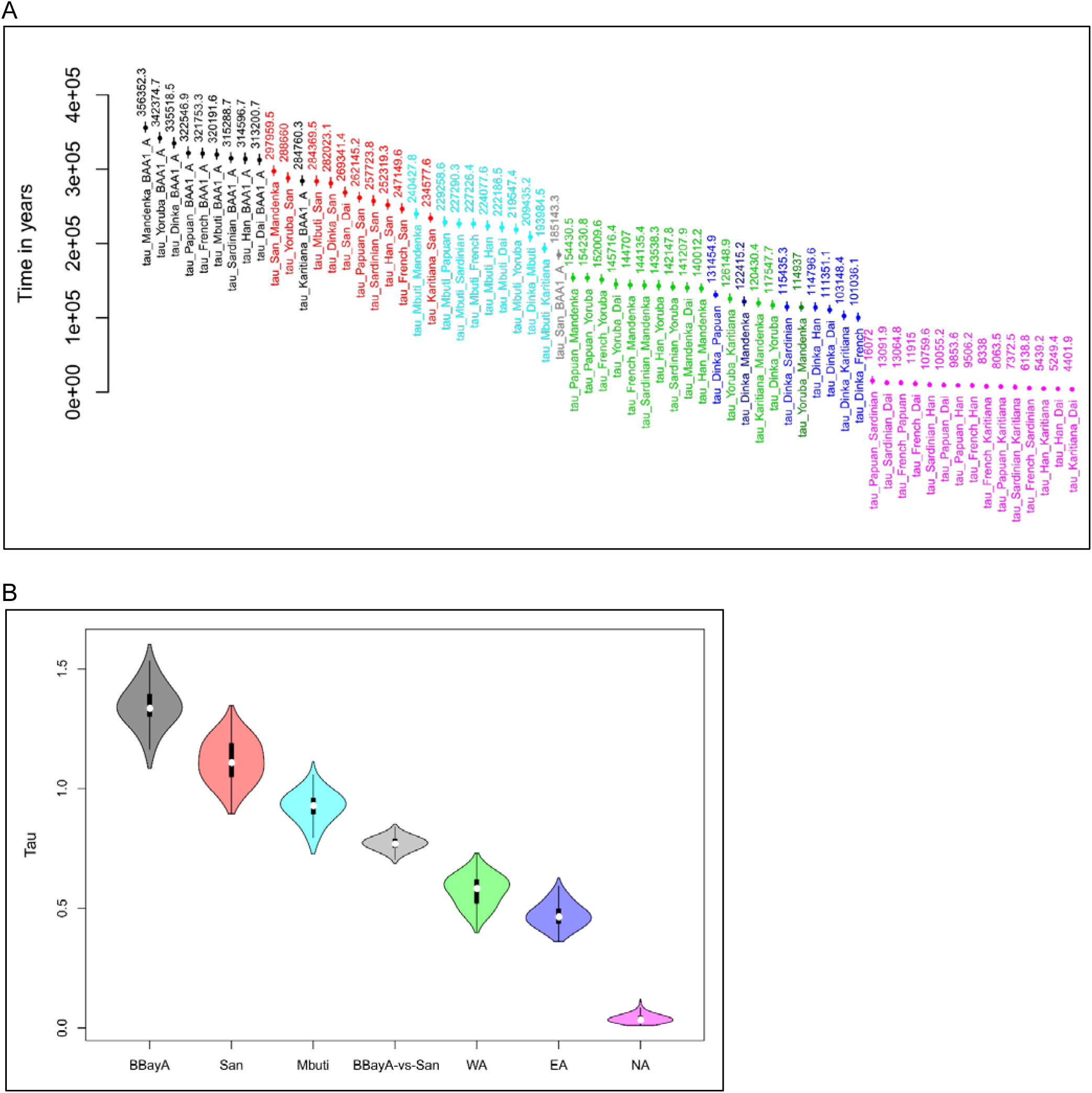
**A)** Means (dots) and standard deviations (bars) of G-PhoCS pairwise population split times, sorted in descending order. Colors are according to the hierarchical split times: Ballito Bay A (BAA) vs. all non-San (Black); San vs. all non-San (Red); Mbuti vs. all non-San (Turquoise); Ballito Bay A vs. San (Gray), West Africans (Mandenka and Yoruba) vs. non-Africans and East Africans (Green); East Africans vs. non-Africans (Blue); pairwise non-Africans (Pink). **B)** Same hierarchical split times shown as violin plots. Y-axis: Tau (Time in generations = Tau / (10,000 × mutation rate)). X-Axis: Populations. West Africans (WA), East Africans (EA), and non-Africans (NA). Means and standard deviations of these grouped split times are summarized in Table S8.1

**Extended Data Fig. 6:**
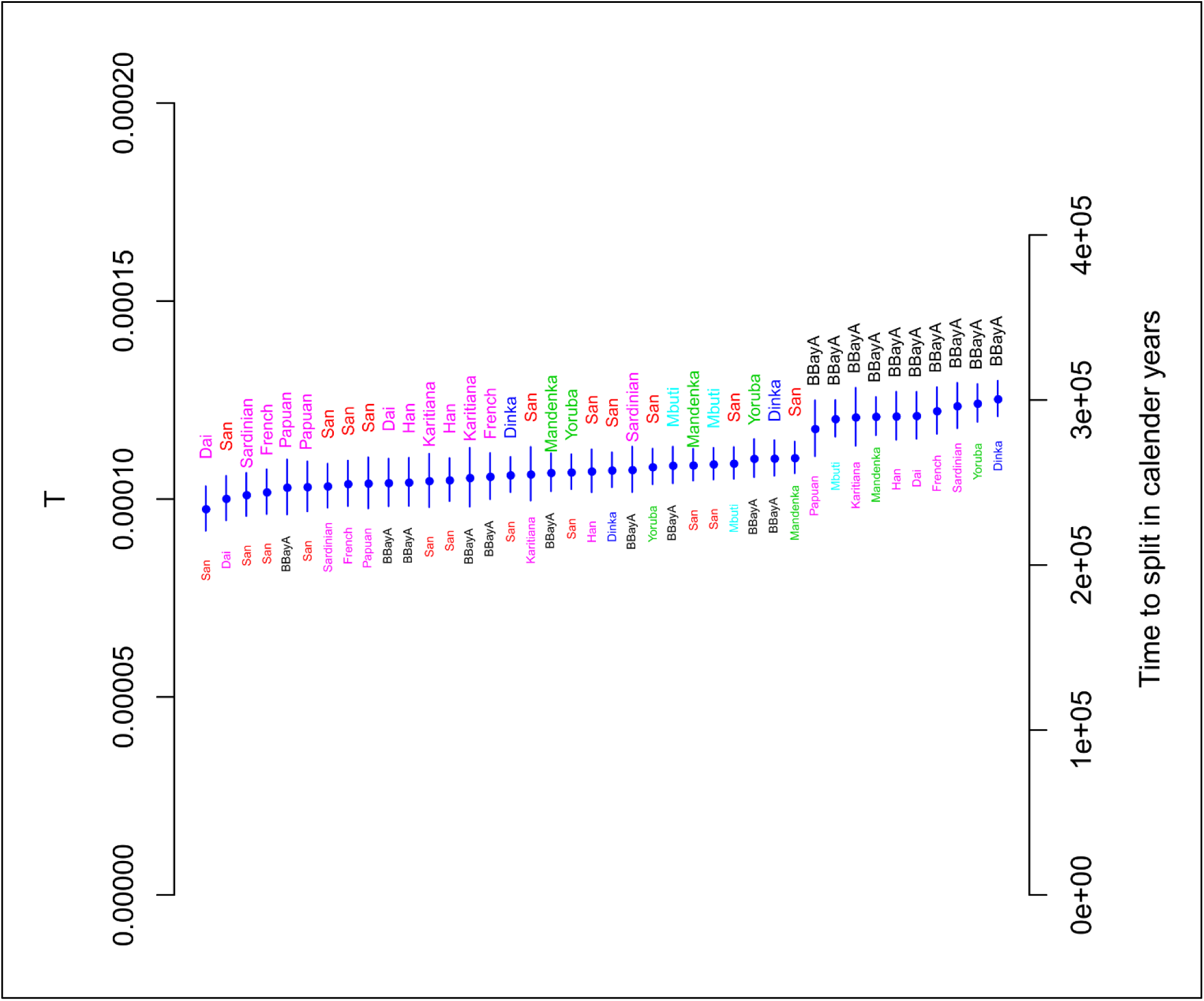
Estimates of split time between pairs of individuals using the TT method. The populations displayed on top and in larger font are focal populations while the populations below in smaller font are the contrasting populations. We assume a mutation rate of 1.25×10^−8^ per site and generation, and a generation time of 30 years to translate the estimated parameter T to time in calendar years.

**Extended Data Fig. 7:**
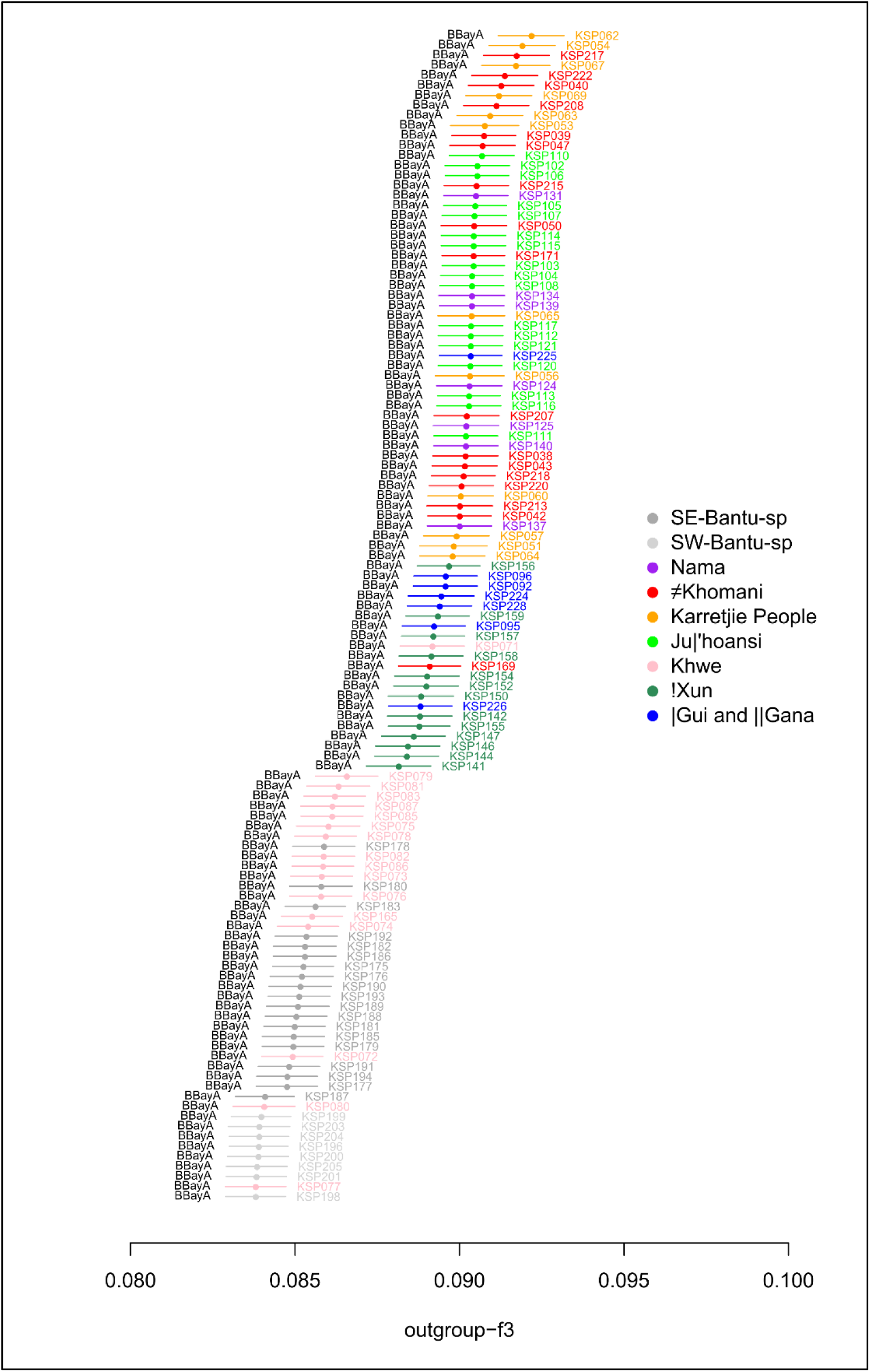
Outgroup-f_3_ between Ballito Bay A and Khoe-San and Bantu-speaker individuals from Schlebusch et al^8^. Ancestral sites are inferred from the genomes of three great apes and are used as outgroup.

**Extended Data Fig. 8:**
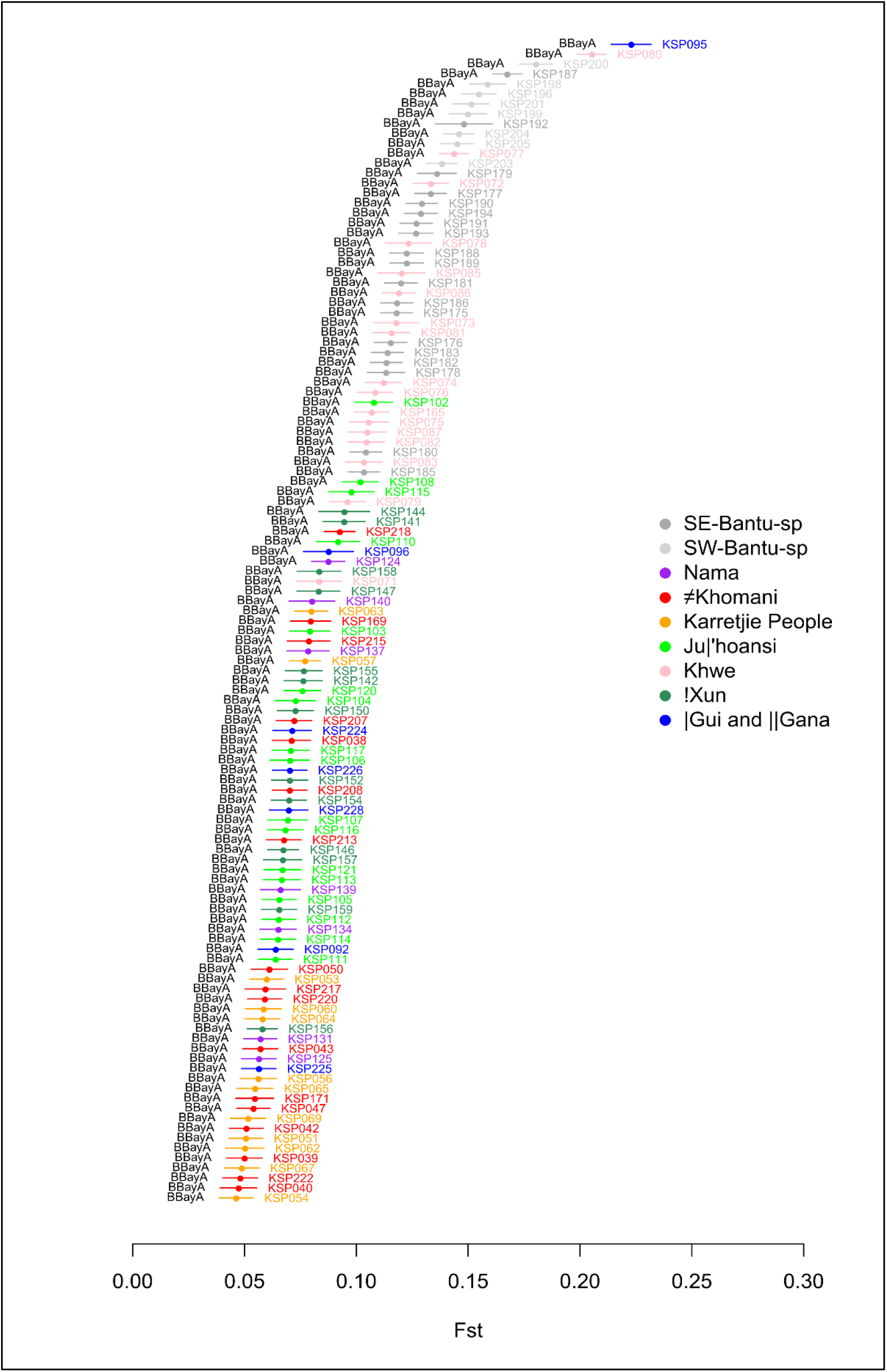
Pairwise F_ST_ between Ballito Bay A and the individuals in Schlebusch et al^8^.

